## Supplementary Figures for "Adolescent neurostimulation of dopamine circuit reverses genetic deficits in frontal cortex function"

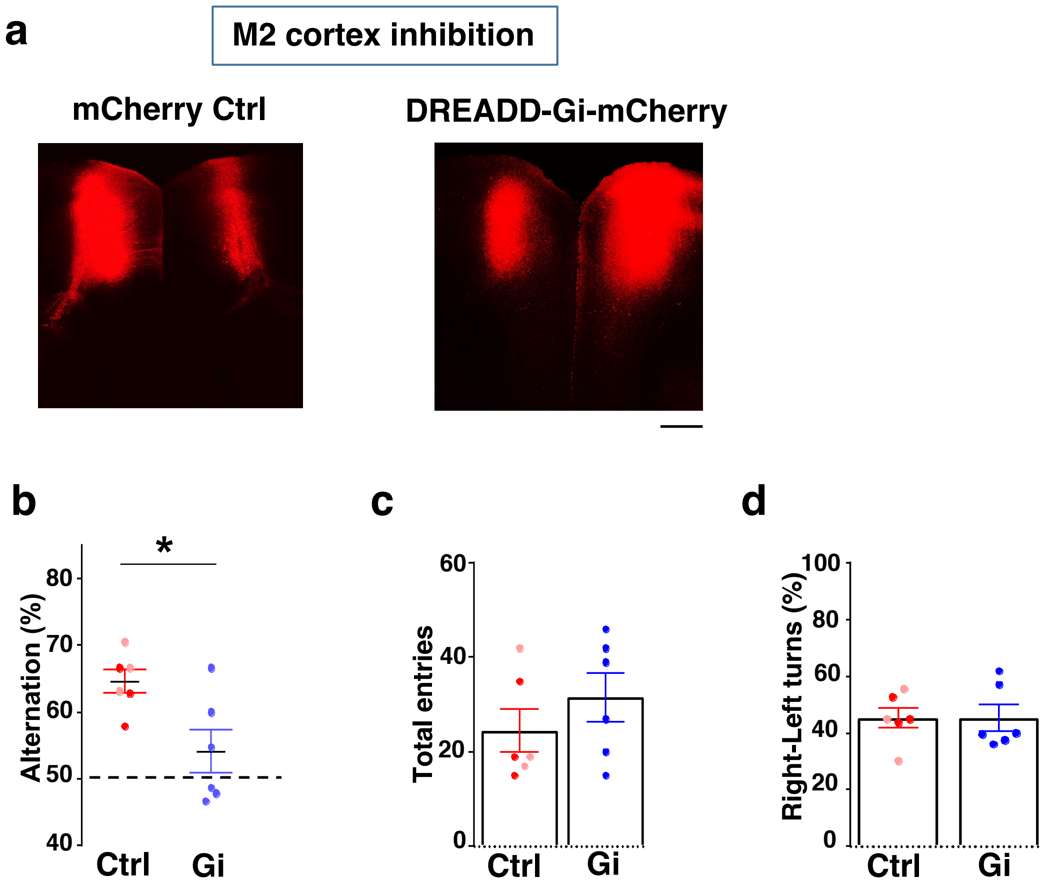
a
M2 cortex inhibition
mCherry Ctrl
DREADD-Gi-mCherry
b
*
Alternation (%)
80
70
60
50
40
Ctrl
Gi
c
Total entries
60
40
20
0
Ctrl
Gi
d
Right-Left turns (%)
100
80
60
40
20
0
Ctrl
Gi

Figure S1

**Fig. S1. Chemogenetic inhibition of M2 frontal cortex reduces alternation in Y-maze.**

(**a**) Confocal image showing expression of control mCherry or DREADD-Gi-mCherry in M2 frontal cortex. Scale bar, 500 $\mu$m. (**b**) Alternation in the Y-maze task showing significant reduction in the Gi animals compared to control animals (*p=0.017, t-test, t(10)=2.845, Gi N=6, Ctrl N=6, including 3 saline-control in Gi labeled mice (pink dots) and 3 mCherry-control with CNO injection (red dots). These two control groups showed no significant difference, and their data were pooled together). (**c**) Total arm entries and (**d**) left-right turns (percentage of right turns shown) are not significantly different. All the error bars indicate SEM.

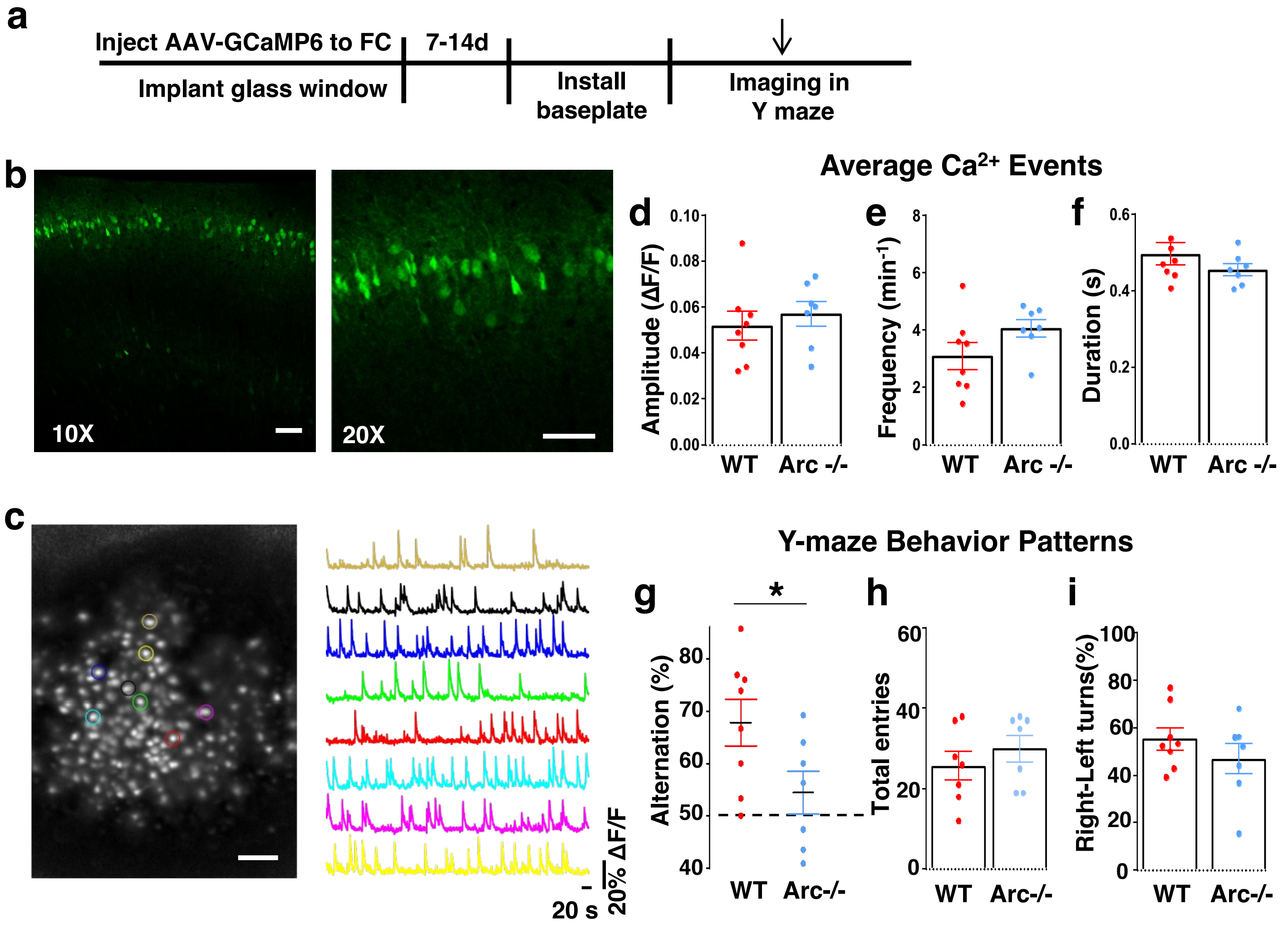


**j**

| | Ca$^{2+}$ Frequency vs. Alternation scores | | Ca$^{2+}$ Frequency vs. Total entries | |
|---|---|---|---|---|
| | $R^2$ | P value | $R^2$ | P value |
| WT (n = 8) | 0.0481 | 0.6019 | 0.0005 | 0.9601 |
| Arc-/- (n = 7) | 0.1816 | 0.3404 | 0.1527 | 0.3861 |
| WT & Arc-/- (n = 15) | 0.0367 | 0.4938 | 0.1270 | 0.1924 |

**Figure S2**

**Fig. S2. Imaging cortical neuronal activity in Y-maze alternation task with a miniaturized head-attached microscope.**

(**a**) Schematic drawing of the outline of experimental procedures. (**b**) Confocal fluorescence images of coronal brain sections showing superficial layer cortical neurons that are labeled by infusion of AAV9-GCaMP6 virus from the pial surface. Left:10X, Right: 20X. Scale bar 100 $\mu$m. (**c**) Left: A representative standard deviation projection image of 15000 frames collected over 8 minutes in the M2 frontal cortex of a freely moving mouse in a Y-maze. Right: The $\Delta F/F$ activity traces of labeled cortical neurons matched in color with the left image. Scale bar 100 $\mu$m. (**d-f**) Quantifications of average calcium transient amplitude **d**, frequency **e**, and duration **f,** in *WT* and *Arc*-/- mice during Y-maze exploration. Each dot on the plot represented the cell average from one mouse. No significant difference was detected. **d**, p=0.548, t-test, t(13)=0.617; **e**, p=0.117, t-test, t(13)=1.679; **f**, p=0.264, t-test, t(13)=-1.168. (**g**) Y-maze alternation percentage for animals in the miniscope imaging experiments, showing reduction in *Arc*-/- animals compared to *WT* (*p=0.047, t-test, t(13)=2.196, *WT* N=8, *Arc*-/- N=7 mice, both groups passed Shapiro-Wilk normality test at alpha=0.05). (**h**) Total entries. (**i**) left-right turns (percentage of right turns shown) are not significantly different. **h**, p=0.69, t-test, t(13)=0.408; **i**, p=0.304, t-test, t(13)=-1.07. All the error bars indicate SEM. (**j**) Pearson pairwise correlation between the frequency of calcium events and the alternation scores or total entries in WT, Arc-/-, and both genotype groups. No significant correlation was found.

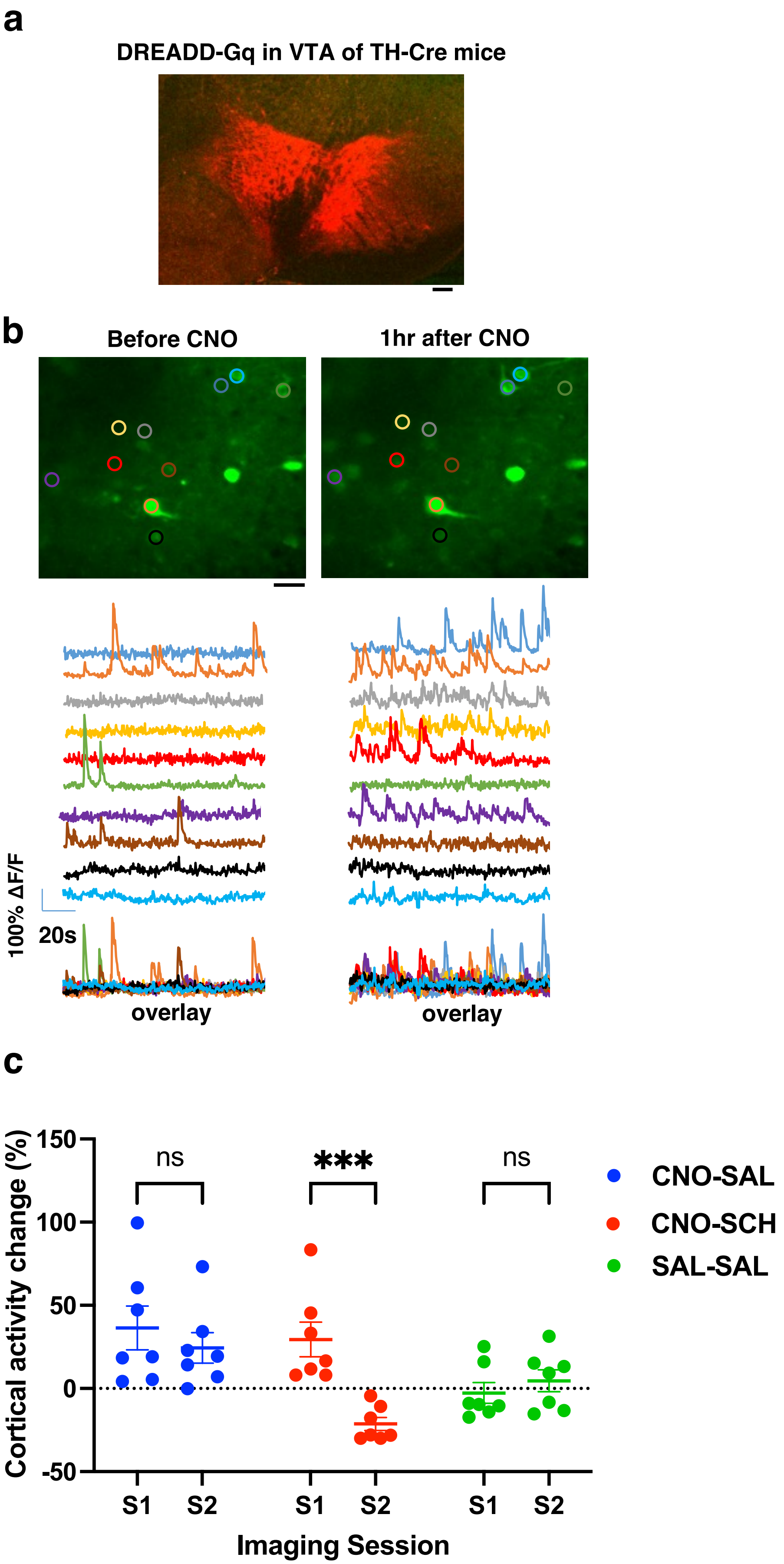

a
DREADD-Gq in VTA of TH-Cre mice
b
Before CNO
1hr after CNO
100% ΔF/F
20s
overlay
overlay
c
Cortical activity change (%)
150
100
50
0
-50
ns
***
ns
S1
S2
S1
S2
S1
S2
Imaging Session
CNO-SAL
CNO-SCH
SAL-SAL


Figure S3

**Fig. S3. VTA expression of DREADD-Gq and validation of CNO induced cortical activation.**

(**a**) Example confocal image of DREADD-Gq-mCherry labeling in the midbrain. Scale bar, 100$\mu$m. (**b**) Top, example images showing standard-deviation projection of *in vivo* two-photon calcium activity movies in the M2 frontal cortex. Bottom, example traces of spontaneous cortical activity in a representative sample of individual cells before and 1hr after CNO injection in a DREADD-Gq animal. Scale bar, 20$\mu$m. (**c**) Saline injection in DREADD-Gq expressing mice did not alter M2 neural activity, and CNO-induced increase in M2 neural activity was suppressed by D1 antagonist SCH23390 (1 mg/kg, i.p.). Two injections (CNO-SAL, CNO-SCH, or SAL-SAL) were done in each animal. Three imaging sessions were conducted: before any injection, 1 hr after the first injection to allow CNO to take effect, and 30 min after the second injection to allow SCH to take effect. Cortical activity is summarized by the standard deviation (SD) of the spontaneous activity traces. Activity changes are calculated as $(SD-SD_0)/SD_0$, where $SD_0$ is before any drug or saline injection, and SD is after CNO, Saline, or SCH23390 injection (2 Way Repeated Measures ANOVA, group x treatment, $F(2,18) = 8.9$, $p = 0.0020$; Sidak's multiple comparison: CNO followed by SAL, $p = 0.5667$; CNO followed by SCH, $p = 0.0002$ ***; SAL followed by SAL, $p = 0.8492$; N = 7 mice for each group).

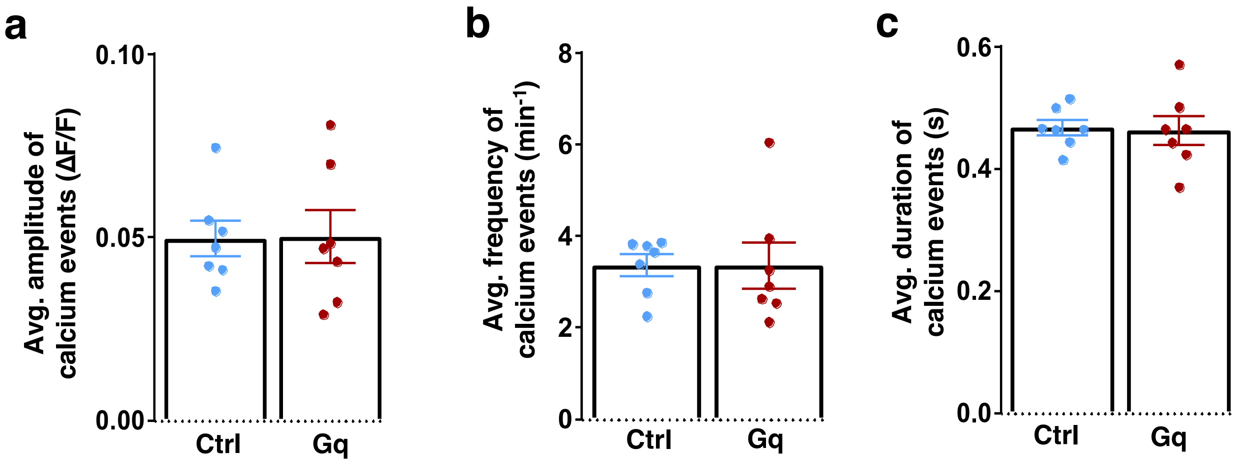
a
Avg. amplitude of
calcium events (ΔF/F)
0.10
0.05
0.00
Ctrl
Gq
b
Avg. frequency of
calcium events (min-1)
8
6
4
2
0
Ctrl
Gq
c
Avg. duration of
calcium events (s)
0.6
0.4
0.2
0.0
Ctrl
Gq

**Figure S4**

**Fig. S4. Average activity of M2 neurons during Y-maze exploration.**

(**a-c**) Quantifications of calcium transient amplitude (**a**), frequency (**b**), and duration (**c**)**,** in *Arc*-/-; mCherry-Ctrl and *Arc*-/-;DREADD-Gq mice during Y-maze exploration show no significant difference. **a**, $p=0.927$, t-test, $t(12)=0.094$; **b**, $p=0.972$, t-test, $t(12)=-0.036$; **c,** $p=0.853$, t-test, $t(12)=-0.189$. Each dot on the plot represented the cell average from one mouse. All the error bars indicate SEM.

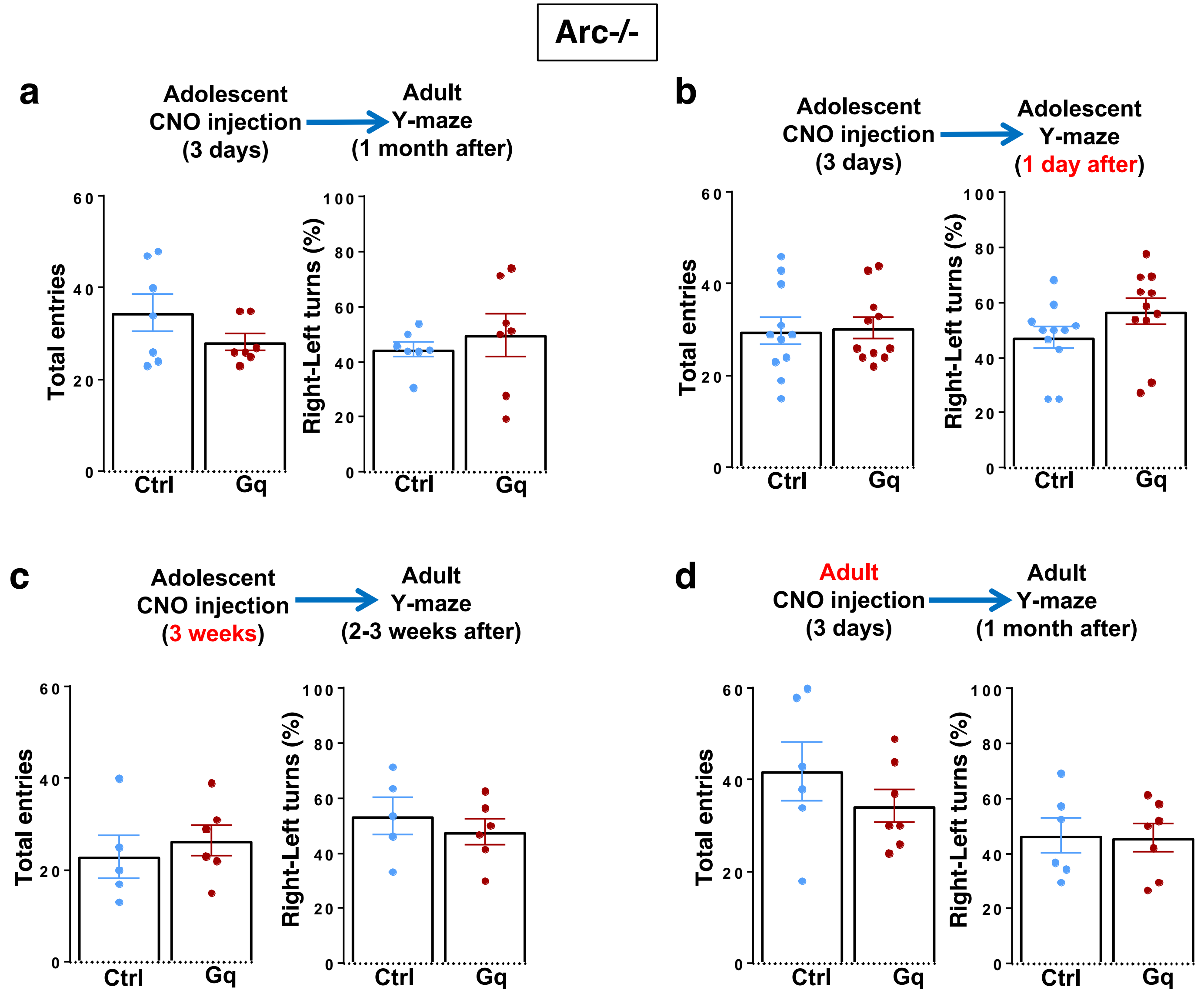

Arc-/-
a
Adolescent CNO injection (3 days)
Adult Y-maze (1 month after)
Total entries
Right-Left turns (%)
Ctrl
Gq
b
Adolescent CNO injection (3 days)
Adolescent Y-maze (1 day after)
Total entries
Right-Left turns (%)
Ctrl
Gq
c
Adolescent CNO injection (3 weeks)
Adult Y-maze (2-3 weeks after)
Total entries
Right-Left turns (%)
Ctrl
Gq
d
Adult CNO injection (3 days)
Adult Y-maze (1 month after)
Total entries
Right-Left turns (%)
Ctrl
Gq


**Figure S5**

**Fig. S5. Behavioral characterization of *Arc*-/- mice under different chemogenetic stimulation conditions.**

(**a**) Top, diagram showing procedures for the stimulation of midbrain dopamine neurons and Y-maze testing in *Arc*-/-;*TH-Cre* mice labeled with DREADD-Gq or mCherry-Ctrl viruses. Total entries and left-right turns (percentage of right turns shown) in the Y-maze are not significantly different between mCherry-Ctrl and DREADD-Gq animals at adulthood after 3-day adolescent CNO stimulation. (**b**) Total entries and left-right turns (percentage of right turns shown) in the Y-maze are not significantly different between mCherry-Ctrl and DREADD-Gq animals 1 day after 3-day adolescent CNO stimulation. (**c**) Total entries and left-right turns (percentage of right turns shown) in the Y-maze are not significantly different between mCherry-Ctrl and DREADD-Gq animals at adulthood after 3-week adolescent CNO stimulation. (**d**) Total entries and left-right turns (percentage of right turns shown) in the Y-maze are not significantly different between mCherry-Ctrl and DREADD-Gq animals after 3-day adult CNO stimulation. All the error bars indicate SEM.

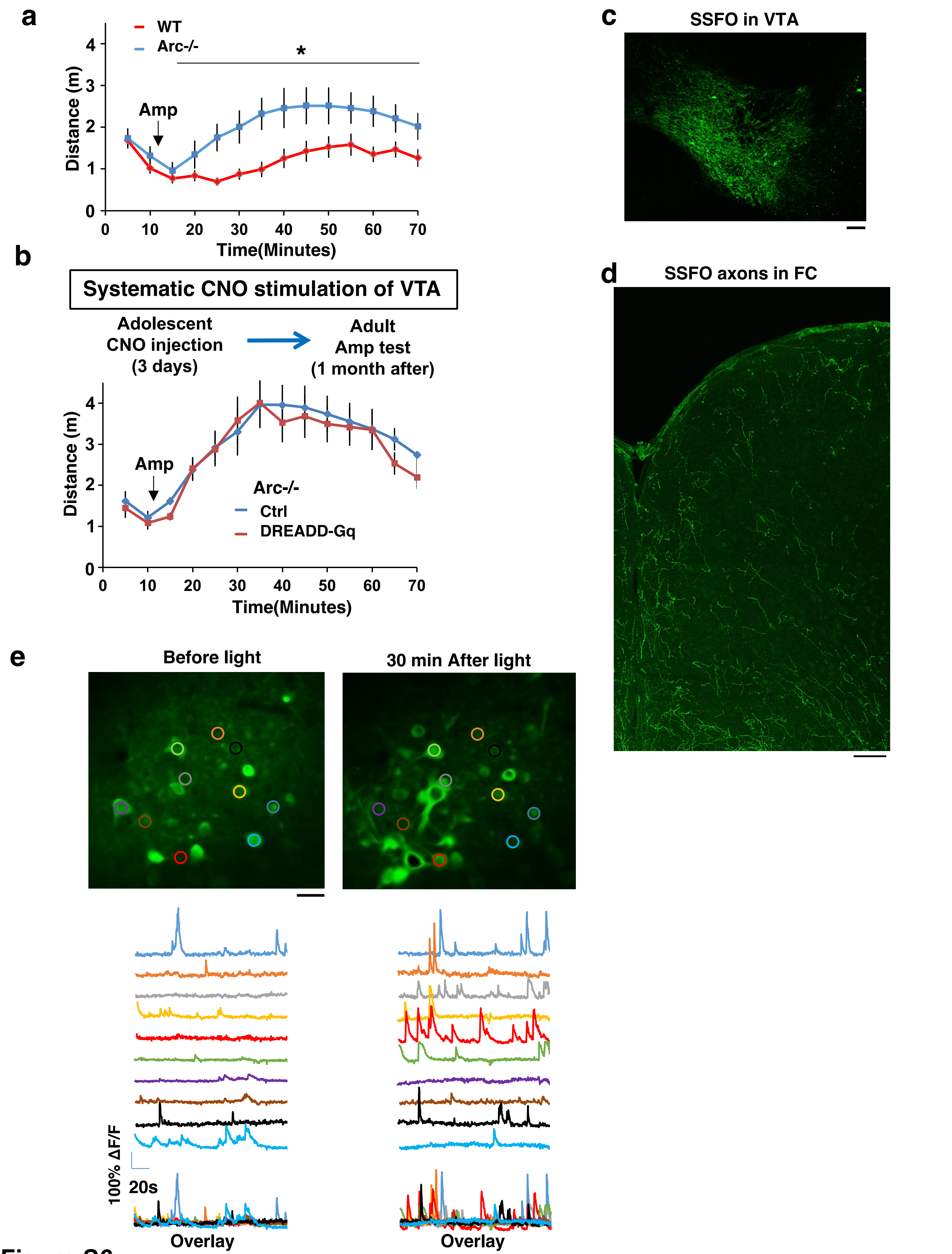
a
WT
Arc-/-
*
Amp
Distance (m)
Time(Minutes)
b
Systematic CNO stimulation of VTA
Adolescent CNO injection (3 days)
Adult Amp test (1 month after)
Amp
Arc-/-
Ctrl
DREADD-Gq
Distance (m)
Time(Minutes)
c
SSFO in VTA
d
SSFO axons in FC
e
Before light
30 min After light
100% ΔF/F
20s
Overlay
Overlay

Figure S6

**Fig. S6. Amphetamine hyper-reactivity in *Arc-/-* mice and SSFO expression and validation.**

(**a**) Amphetamine induced locomotion is significantly increased in *Arc*-/- animals compared to *WT*. ($F(1,14)=9.5$, *$p=0.032$, Two-way ANOVA, N=8 mice per group). (**b**) Amphetamine induced locomotion at adulthood is not significantly different between *Arc-/-;TH-Cre* mice labeled with mCherry-Ctrl or DREADD-Gq and treated with CNO for 3 days in adolescence. All the error bars indicate SEM. (**c**) Confocal image of coronal midbrain section showing SSFO expression in the VTA. Scale bar, 100 $\mu$m. (**d**) Confocal image of coronal cortical section showing SSFO expression in the frontal dopaminergic axons projecting from the VTA. Scale bar, 20 $\mu$m. (**e**) Top, example images showing standard-deviation projection of *in vivo* two-photon calcium activity movies in the M2 frontal cortex. Bottom, example traces of spontaneous cortical activity in a representative sample of individual cells before and 30 min after light stimulation in a SSFO labeled animal. Scale bar, 20$\mu$m.

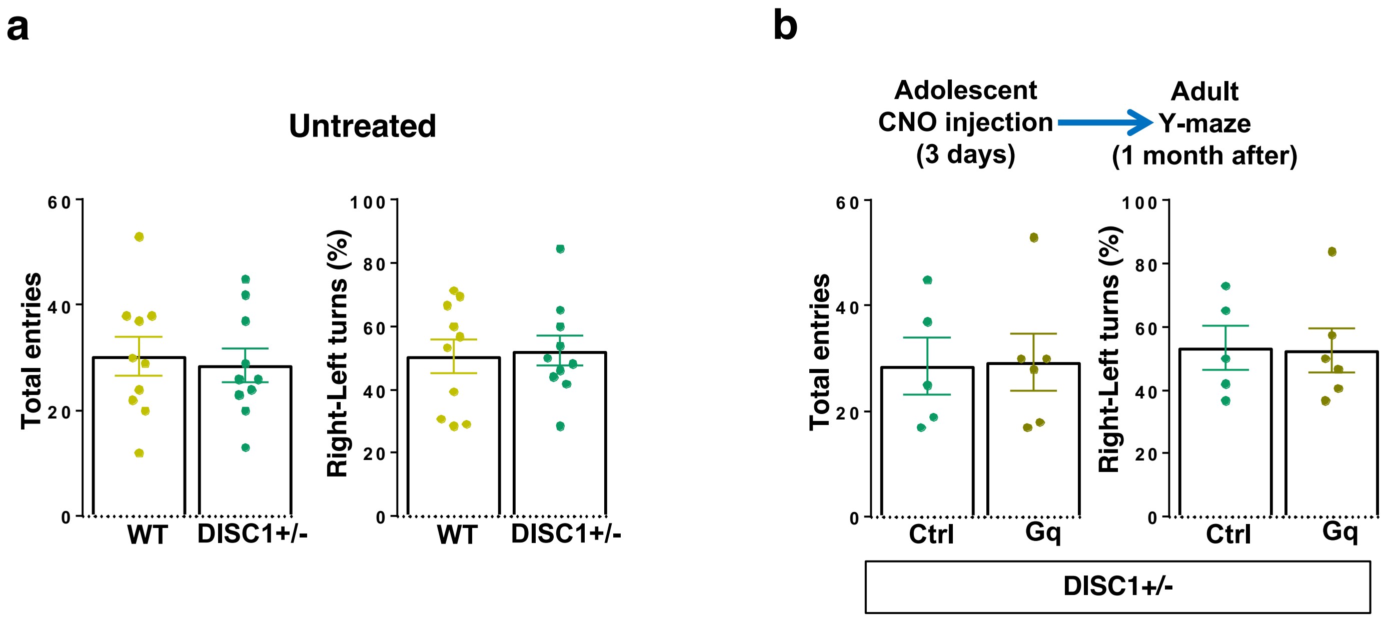
a
Untreated
Total entries
Right-Left turns (%)
WT
DISC1+/-
b
Adolescent
CNO injection
(3 days)
Adult
Y-maze
(1 month after)
Ctrl
Gq
DISC1+/-

Figure S7

**Fig. S7. Characterization of Y-maze behavior in *DISC1+/-* mice.**

(**a**) Total entries and left-right turns (percentage of right turns shown) in the Y-maze are not significantly different between *DISC1*+/- and *WT* animals. (**b**) Top, diagram showing procedures for the stimulation of midbrain dopamine neurons and Y-maze testing in *DISC1*+/-;*TH-Cre* mice labeled with DREADD-Gq or mCherry-Ctrl viruses. Total entries and left-right turns (percentage of right turns shown) in the Y-maze are not significantly different between mCherry-Ctrl and DREADD-Gq animals at adulthood after 3-day adolescent CNO stimulation. All the error bars indicate SEM.

**Supplementary Movie S1. Frontal cortical neuronal ensemble activity in freely moving mice during Y-maze exploration.**

Simultaneously recorded behavioral video of a mouse exploring a Y-maze (left) and calcium activity ($\Delta F/F$) movie of M2 cortical neurons imaged through a head-attached miniaturized microscope (right).
